# Epigenetic Reshaping through Damage: Promoting Cell Fate Transition by BrdU and IdU Incorporation

**DOI:** 10.1101/2023.08.28.555083

**Authors:** Chuang Li, Shuyan Chen, Xiaoduo Xu, Tongxing Guan, Anchun Xu, Chen Li, Shengyong Yu, Haokaifeng Wu, Duanqing Pei, Jing Liu

**Affiliations:** School of Life Sciences, University of Science and Technology of China, Hefei 230026, China; Center for Cell Lineage and Development, Guangzhou Institutes of Biomedicine and Health, Chinese Academy of Sciences, Guangzhou 510530, China; CAS Key Laboratory of Regenerative Biology, Guangdong Provincial Key Laboratory of Stem Cell and Regenerative Medicine, Guangzhou Institutes of Biomedicine and Health, Chinese Academy of Sciences. Guangzhou 510530, China; Centre for Regenerative Medicine and Health, Hong Kong Institute of Science & Innovation, Chinese Academy of Sciences, Hong Kong SAR, P. R. China; University of Chinese Academy of Science, Beijing, 100049, P. R. China; Key Laboratory of Biological Targeting Diagnosis, Therapy and Rehabilitation of Guangdong Higher Education Institutes, The Fifth Affiliated Hospital of Guangzhou Medical University, Guangzhou 510000, China

## Abstract

Thymidine analogs have long been recognized for their ability to randomly incorporate into DNA. However, their significance in the chemical induction of pluripotency (CIP) remains unclear. Here, we investigated the impact of BrdU/IdU incorporation on the transition of cell fate through DNA damage repair (DDR). Our findings reveal a substantial upregulation of reprogramming mediator gene H3K27ac and H3K9ac, as well as global DNA demethylation in response to DDR. This process creates a hypomethylated environment that promotes cell fate transition. We term this mechanism as Epigenetic Reshaping through Damage (ERD). Overall, our study sheds light on the dynamic interplay between thymidine analogs, DDR, and epigenetic modifications, providing valuable insights into the mechanisms underlying cell fate transition.

## Introduction

Thymidine analogs are often considered to incorporate randomly into DNA sequences, introducing inherent unpredictability and randomness in their mechanisms of action(*1*). Despite this inherent nature, they play a critical role in the precise regulation of cell fate(*2–7*). In a study conducted by Xie et al., the utilization of BrdU, widely employed to label proliferating cells in vivo, has demonstrated its significant potential in facilitating chemical induction of pluripotency(*3*). The study by Cao et al., sheds light on how BrdU can impact the reorganization of nuclear architecture and influence the cell fate decisions(*5*). IdU, which is also a thymidine analog was reported to cause stochastic fluctuations in gene expression to facilitate cellular reprogramming(*8*).

However, the precise mechanisms through which BrdU/IdU facilitates cell reprogramming are not yet fully understood. BrdU/IdU may be involved in cell reprogramming through mechanisms beyond transcriptional fluctuations. This could include the impact of thymidine analogs on DNA structure, repair mechanisms, or other cellular processes, providing a potential shortcut for cells to overcome reprogramming barriers. As a result, thymidine analogs may play a more crucial and multifaceted role in regulating cell fate, extending beyond the induction of transcriptional fluctuations alone. This study links DNA damage repair with genome, epigenetics and cell fate regulation, providing us with new understanding and helping to advance the field of cell fate decision and regenerative medicine.

## Results

### BrdU and IdU: Essential in overcoming two barriers during chemical induction of pluripotency

To determine the optimal duration of BrdU treatment for CIP, a series of time windows were optimized and observed for their impact on CIP (Figure 1A). Two main barriers were found in the CIP reprogramming process, and BrdU was found to play a crucial role in overcoming these barriers. BrdU dropout experiments showed that dropout of BrdU at barrier I resulted in a failure to form colonies at Day 22, while dropout at barrier II between Day 12-22 resulted in a failure to form GFP^+^ colonies at Day 40 (Figure 1B). Meanwhile, we have established a time gradient for each barrier (Figure S1A), and breaking through the barrier requires at least 6 days of BrdU incorporation, which must be added within the first 4 days (Figure S1B). Increasing the duration of BrdU treatment led to an increase in the number of GFP^+^ colonies, with a 0-22 days treatment resulting in over 240 GFP^+^ colonies at Day 40 from a starting population of 20,000 cells, achieving an efficiency of 1.2%. In contrast, a 0-12 days treatment only resulted in about 20 GFP^+^ colonies (Figure 1C and S1C). Furthermore, we attempted to use three base analogs, namely IdU, EdU, and 5Aza, but only IdU was found to be capable of replacing the function of BrdU during CIP reprogramming (Figure 1D and S1D). The combination of both thymidine analogs can increase the selectivity of cells, so that only cells that have been correctly reprogrammed can survive; hence accelerated the reprogramming process (Figure 1D,E and S1E). Thus, it can be concluded that BrdU and IdU are essential in overcoming two main barriers during CIP.

**Fig. 1.**
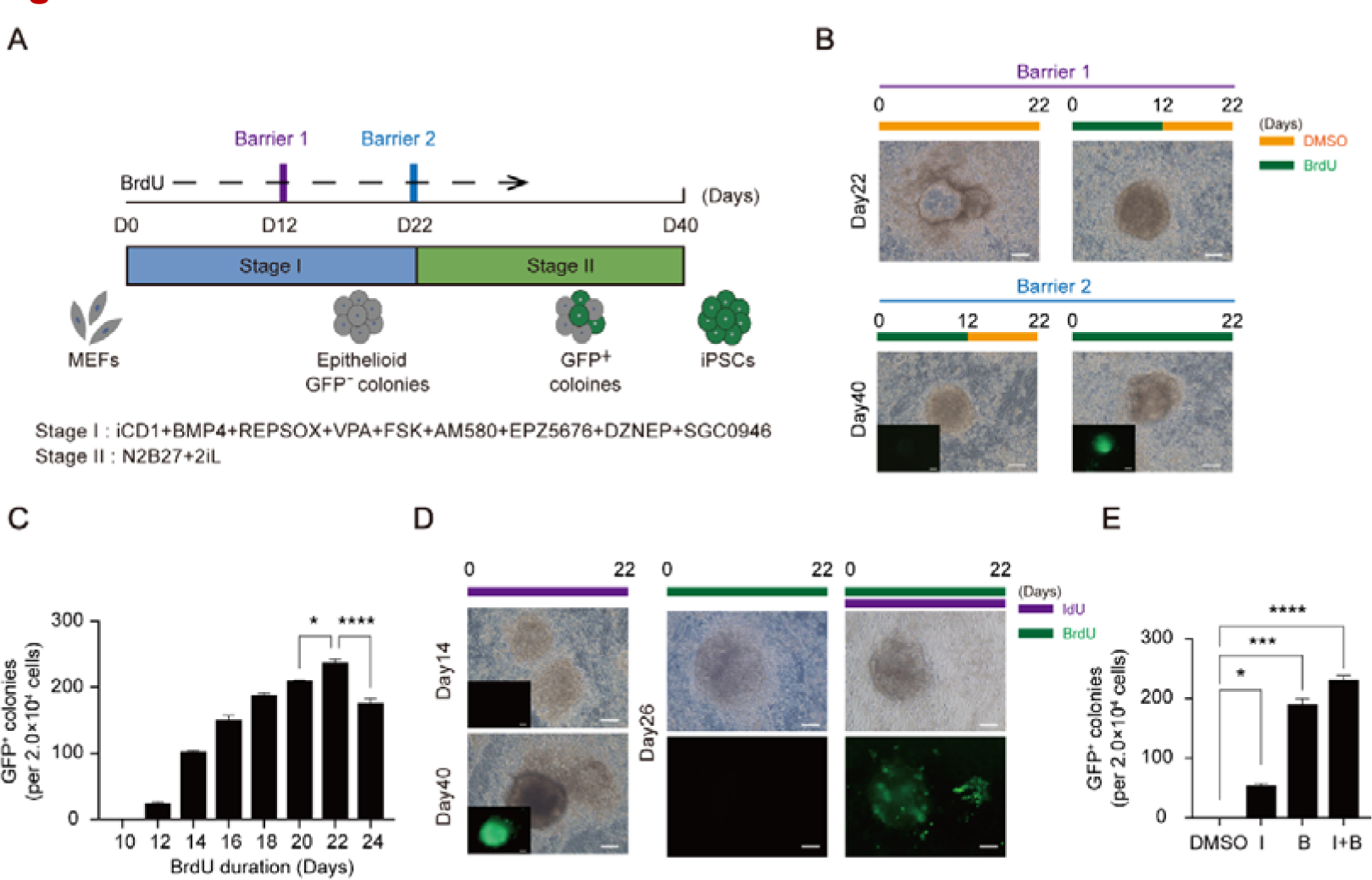
BrdU and IdU: Essential in overcoming two barriers during chemical induction of pluripotency. (**A**) Schematic diagram of the induction of CiPSCs from MEFs. (**B**) Morphological changes at two distinct barriers during induction of CiPSCs. Scale bar 100 µ m. (**C**) Number of Oct4-GFP+CiPSC colonies generated under indicated conditions. (**D**) Morphological changes at distinct time points during induction of CiPSCs treated with BrdU or IdU. Scale bar 100µm. (**E**) Number of Oct4-GFP+CiPSC colonies generated under different treatment conditions.

### Gene expression dynamics and chromatin accessibility dynamics during CIP with or without BrdU/IdU

To identify the two barriers in which BrdU and IdU are involved during CIP reprogramming, RNA-seq and ATAC-seq was performed on MEFs undergoing CIP with or without BrdU, IdU and I+B (IdU+BrdU) at D8, 14, 20, 26 and 40 (Figure 2A). It was found that loss of BrdU and IdU at barrier I resulted in incomplete reprogramming of MEFs to XEN-like cells (Figure 2B). The expression of XEN-like genes such as *Gata4*, *Sox17* and *Aqp8* increased significantly with increasing duration of thymidine analogs treatment and reached a peak at D20. Conversely, dropout of BrdU resulted in a failure to express XEN-like genes (Figure S2A,C), suggesting that MEFs undergoing reprogramming without BrdU /IdU follow a different fate path. Previous studies have reported that CIP goes through an XEN-like intermediate stage(*2*) . Additionally, loss of BrdU and IdU at barrier II resulted in incomplete reprogramming of XEN-like cells to CiPSCs. Pluripotency genes such as *Oct4*, *Esrrb, Tfcp2l1*, *Nanog* and *Sox2* were highly activated in a BrdU/IdU dependent manner during stage 2. In addition, the incorporation of I+B showed a faster transition towards to iPSCs compared to BrdU/IdU treatment at the RNA level (Figure S2B,D).

**Fig. 2.**
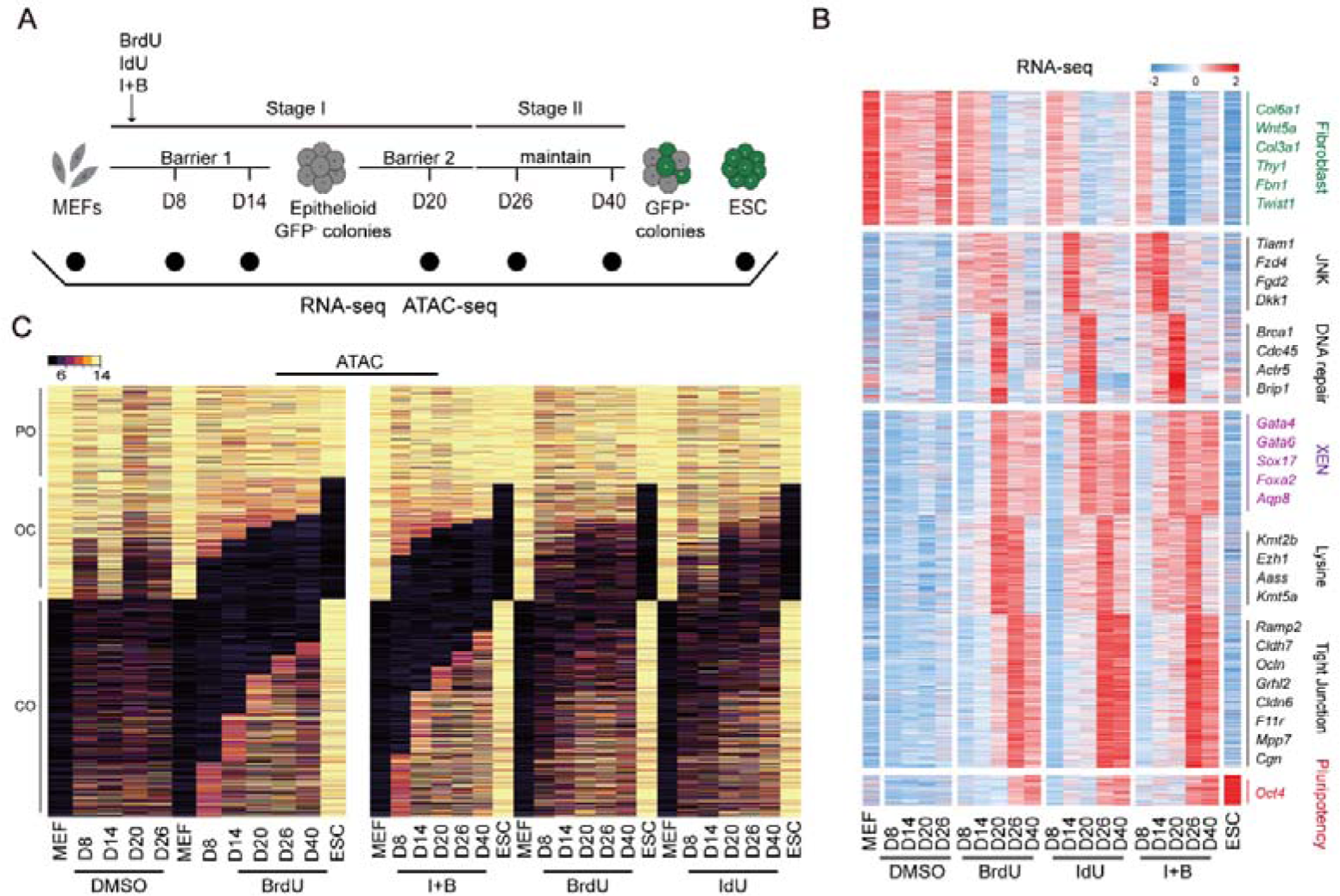
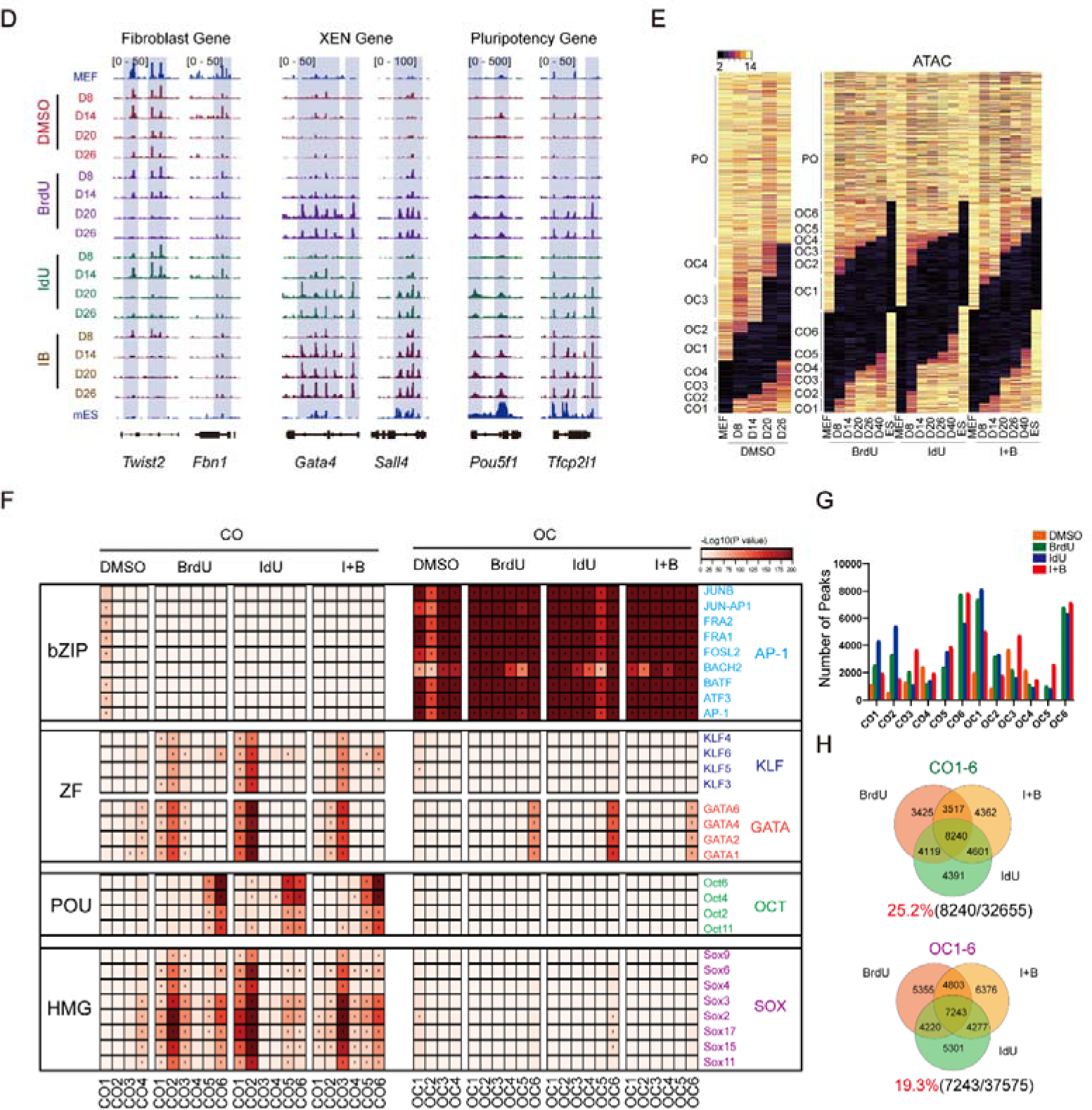
Gene expression dynamics and chromatin accessibility dynamics during CIP with or without BrdU/IdU. (**A**) Schematic diagram of the induction of CiPSCs from MEFs. (**B**) Heatmap of RNA-seq for CIP at D8,14,20 and 26 with or without BrdU or IdU. (**C**)The global chromatin status of CO/OC and PO for CIP at D8,14,20,26 and 40 with or without BrdU or IdU. (**D**) Representative fibroblast genes, XEN genes and pluripotency genes peaks from ATAC-seq for CIP at D8,14,20 and 26 with or without BrdU or IdU. (**E**) The global chromatin status of CO/OC and PO arranged as groups for CIP at D8,14,20,26 and 40 with or without BrdU or IdU. (**F**) TF motifs enriched at least 2 fold for CO/OC loci defined by ATAC-seq peaks with or without BrdU or IdU. TF families are indicated on the right of the heatmap. (**G**) Statistical analysis of number of total peaks in CO1-6/OC1-6. (**H**) Venn diagrams of OC1-6/CO1-6 peaks between samples with or without BrdU or IdU.

The function state of a cell is determined by its genome architecture. To investigate the role of BrdU/IdU in CIP, we mapped chromatin accessibility dynamics (CADs) during CIP with or without BrdU/IdU and found that many loci opened during CIP failed to open without BrdU/IdU. Moreover, detailed analysis of the ATAC-seq datasets revealed many similarities between BrdU and IdU treatments. Chromatin accessibility significantly increased when MEFs undergoing CIP were treated with I+B compared to only BrdU or IdU treatment (Figure 2C). In Figure 2D, we analyzed the loci near fibroblast marker gene *Twist2* and *Fbn1* found that they were more open in DMSO treatment compared to BrdU/IdU treatment. Conversely, loci near XEN marker genes such as *Gata4* and *Sall4* were only open in BrdU/IdU treatment. Loci near the pluripotency marker gene *Oct4* and *Tfcp2l1* were also only open in BrdU/IdU treatment, and the chromatin accessibility was much higher when samples were treated with both I+B.

We compared the peaks at each locus between these samples and classified them into three categories: closed in MEFs but open in ESCs (CO), open in MEFs but closed in ESCs (OC), and permanent open (PO). Further, we divided the CO and CO peaks into subgroups based on the day of reprogramming to illustrate the progression of cellular reprogramming associated CADs, as shown in Figure 2E. To further investigate the molecular mechanism of BrdU/IdU during CIP, we performed motif analysis, as illustrated in Figure 2F. Loci containing motifs for OCT2, 4, 6, and 11 gradually opened, peaking at CO6 when treated with BrdU/IdU. Loci with motifs for GATA1, 2, 4, 6, KLF3-6, and SOX2-4, 6, 9, 10, 15, 17 opened from CO1-6 when treated with BrdU/IdU. However, the TF motifs profile was similar with or without BrdU/IdU in OC loci, indicating that BrdU/IdU treatment tended to open up loci that are enriched with motifs binding to TFs from the GATA, KLF, and SOX families.

We then statistically analyzed the number of total peaks in CO1-6/OC1-6, respectively, and presented the results in the form of a Venn diagram, as shown in Figure 2G,H. The diagram showed that BrdU/IdU/I+B shared 25.2% (8240/32655) in CO peaks, but only 19.3% (7243/37575) in OC peaks. Figure S2E illustrates that a larger fraction of peaks in CO are located in the promoter regions. In conclusion, the function of BrdU/IdU is more biased towards opening chromatin and activating gene expression.

### Incorporation of BrdU/IdU leads to DNA damage repair

According to the RNA-seq analysis we obtained in Figure 2, we then performed Venn plots. Interestingly, Venn plots for overlapping genes in the upregulated groups between BrdU, IdU, and I+B versus the control (without thymidine analogs) showed that DNA repair-related GO functions were highly enriched when MEFs were treated with BrdU/IdU during CIP (Figure 3A,B). Specifically, DNA repair-related gene expression was highly upregulated in all three groups. The main repair pathways are base excision repair (BER), nucleotide excision repair (NER), mismatch repair (MMR), homologous recombination repair (HRR) and non-homologous end joining repair (NHEJ). Among which HRR is the most dominant repair pathway. Additionally, a combination of both thymidine analogs (I+B) accelerated the DNA repair process, as shown by the early peak of DNA repair gene expression at D8 in this group compared to the BrdU or IdU only groups (Figure 3C).

**Fig. 3.**
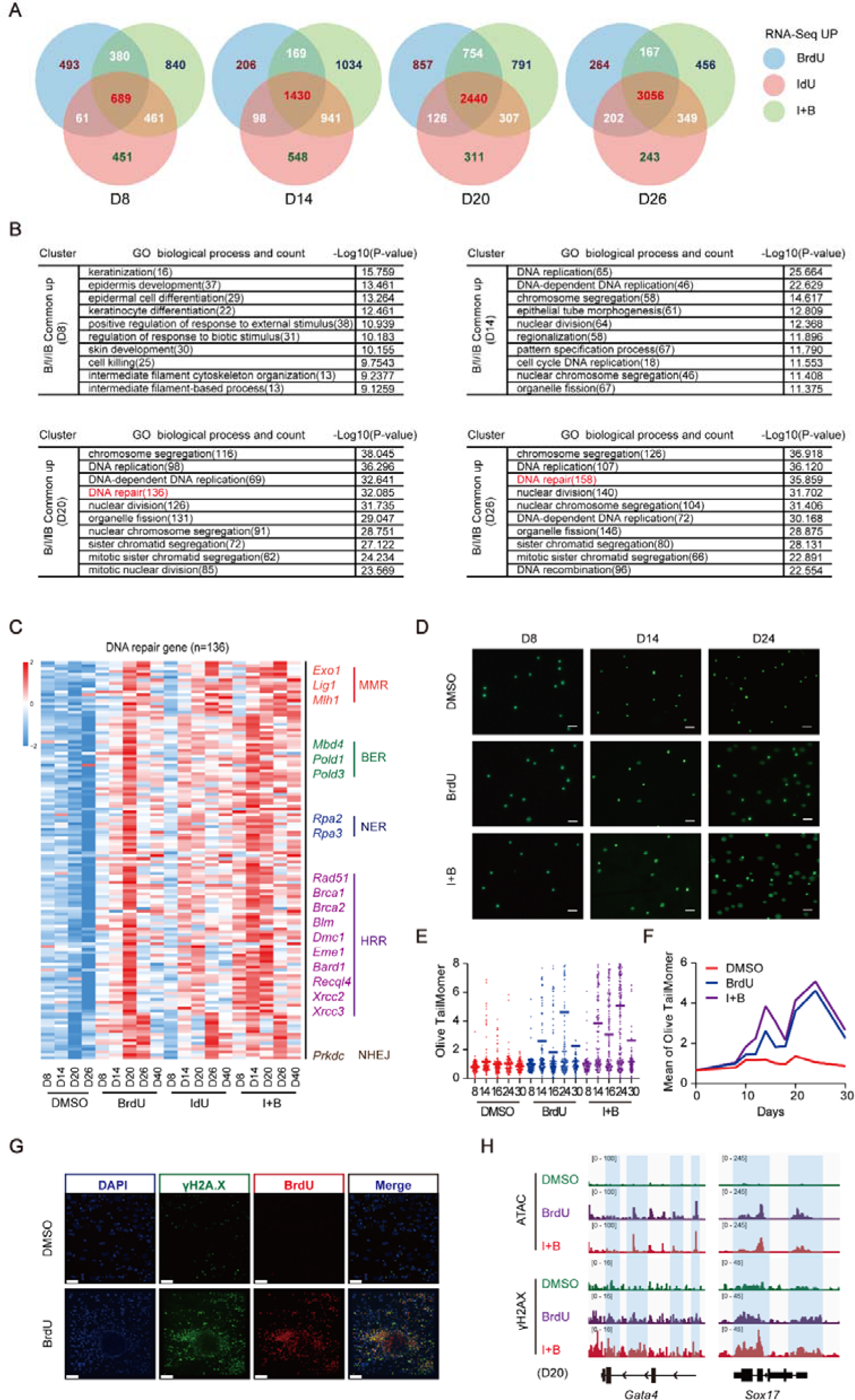
Incorporation of BrdU/IdU leads to DNA damage repair. (**A**) Venn plots for overlapping genes in the upregulated groups between BrdU, IdU, and BrdU+IdU versus the control. (**B**) Enriched GO functions in upregulated groups. (**C**) Heatmap of DNA-repair genes related RNA-seq for CIP at D8,14,20 and 26 with or without BrdU or IdU. (**D**) Alkaline comet assays of samples with or without BrdU or IdU. Scale bar 50µm. (**E**) Statistical analysis of olive tailmomer from (**D**). (**F**) Statistical analysis of mean tailmomer from (**E**). (**G**) Immunofluorescent analysis to observe the spatial colocalization of BrdU (in red) and γ-H2AX (in green). Scale bar 40µm. (**H**) Representative peaks from ATAC-seq aligned with γ-H2AX Cut & tag signals for Gata4 and Sox17 with or without BrdU or IdU.

To verify the relationship between thymidine analogs and DNA damage/repair, alkaline comet assays were performed to compare the olive tail moment difference between samples with or without thymidine analogs treatment (Figure 3D,E). Interestingly, the BrdU treatment showed an increased olive tail moment indicating significant DNA damage. Furthermore, the degree of DNA damage was further promoted by a combination of BrdU and IdU. Figure 3F shows two DNA damage peaks during the CIP process at D14 and D24, which is coherent with the time window of the two main barriers discussed in the previous sections. This evidence illustrates a close association between DNA damage/repair and thymidine analogues assisting CIP through two barriers.

To verify whether BrdU treatment directly causes DNA damage, immunofluorescent analysis was performed to observe the spatial colocalization of BrdU (in red) and γ-H2AX (in green), a novel biomarker for DNA double-strand breaks (Figure 3G). The dropout of BrdU showed a much weaker γ-H2AX signal, leading us to conclude that the accumulation of DNA damage during the CIP process is directly caused by BrdU binding to the DNA. Using CHIP-seq analysis targeting the DNA double-strand damage marker γ-H2AX, a correlation was found between its level and the ATAC-seq for *Gata4* and *Sox17* (Figure 3H). This suggests that the activation of the *Gata*4 and *Sox*17 genes is mediated through the DNA damage repair pathway.

### Mechanism of DDR in promoting somatic cell reprogramming

ATM (Ataxia Telangiectasia Mutated) is a protein kinase that plays a critical role in the cellular response to DNA damage, particularly double-strand breaks (DSBs). When DSBs occur, the MRN complex (Mre11-Rad50-Nbs1) recognizes and binds to the site of break, recruiting ATM. Subsequently, ATM undergoes autophosphorylation, resulting in its full activation. Once activated, ATM phosphorylates several downstream targets involved in cell cycle checkpoint control, DNA repair, and apoptosis. Additionally, ATM activation is sustained by a phosphorylation-acetylation cascade involving c-Abl and TIP60. This cascade helps to maintain ATM activity and promote efficient DNA repair. Overall, ATM serves as a critical player in the cellular response to DNA damage, and its activation and downstream signaling are tightly regulated to ensure proper DNA repair and maintenance of genomic integrity(*9*).

In order to understand how DDR is involved in regulating somatic cell reprogramming, we hypothesize that DDR can upregulate the acetylation levels of genes through the ATM-acetylation mechanism at the site of DNA damage, thereby regulating gene expression, as shown in Figure 4A. Firstly, we found that inhibiting the ATM signaling pathway with an ATM inhibitor prevented BrdU from functioning (Figure 4B,C). Furthermore, we found that *Gata4*, *Sox17* and *Sall4*’s H3KC27ac and H3K9ac were significantly upregulated under the treatment of BrdU and I+B (Figure 4D). To further verify this hypothesis, we increased the concentration of VPA (a histone deacetylase inhibitor) to further promote acetylation accumulation. We found that this significantly accelerated the process of chemical reprogramming, and its acceleration effect depended on the addition of BrdU (Figure 4E,F).

**Fig. 4.**
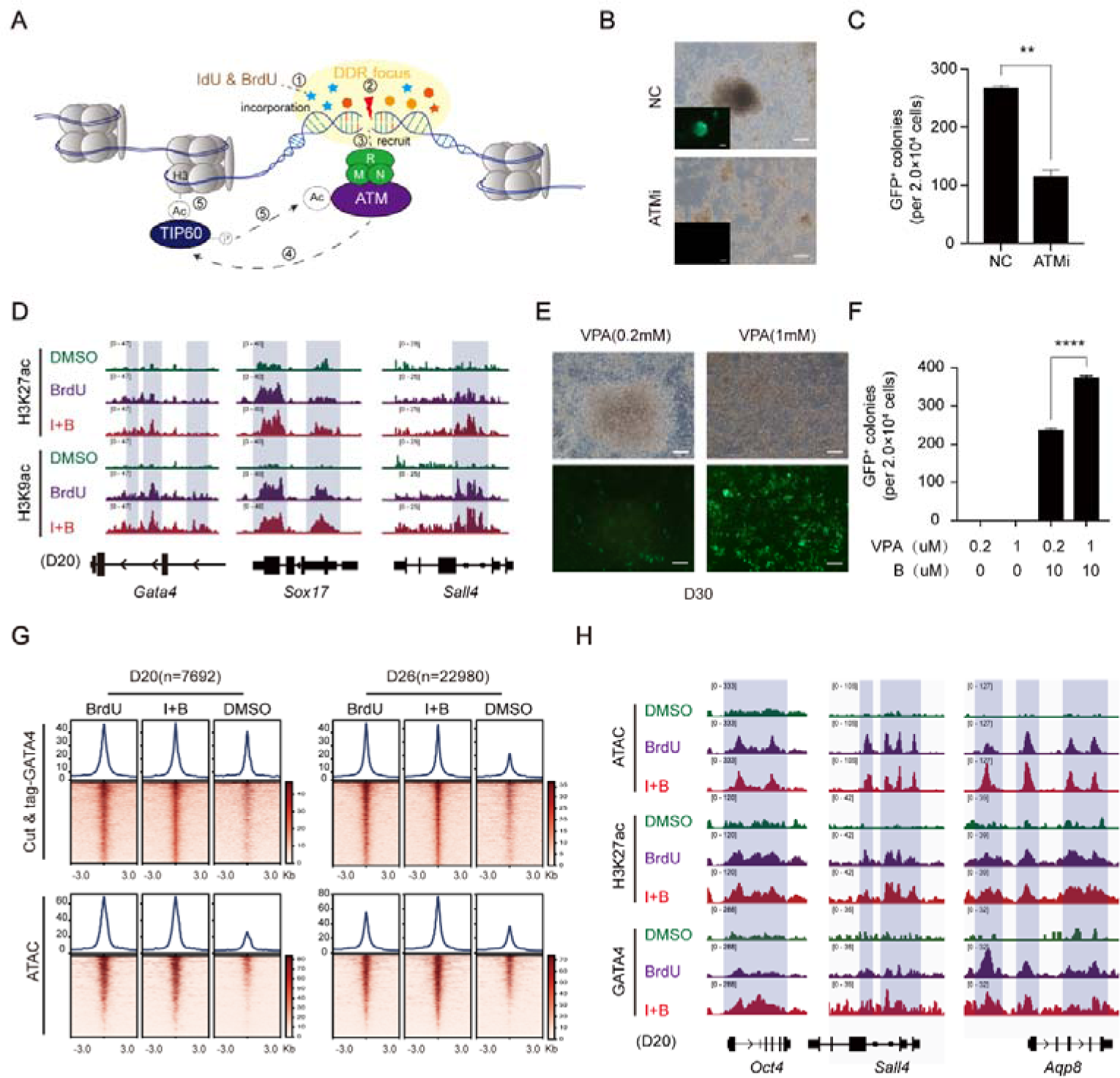
Mechanism of DDR in promoting somatic cell reprogramming. (**A**) A model for DNA damage repair (DDR) mechanism during BrdU/IdU incorporation. (**B**) Morphological changes during induction of CiPSCs treated with or without ATM inhibitor. Scale bar 100µm. (**C**) Number of Oct4-GFP+CiPSC colonies generated under different treatment conditions. (**D**) Representative Gata4, Sox17 and Sall4 peaks from H3K27ac Cut & tag aligned with H3K9ac Cut & tag signals for CIP at D20 with or without BrdU or IdU. (**E**) Morphological changes during induction of CiPSCs treated with different concentration of VPA. Scale bar 100µm. (**F**) Number of Oct4-GFP+CiPSC colonies generated under different treatment conditions. (**G**) GATA4 Cut & tag analysis and ATAC-seq analysis for CIP at D20 and D26 under different treatment conditions. (**H**) Representative Oct4, Sall4 and Aqp8 peaks from ATAC-seq aligned with H3K27ac Cut & tag and GATA4 Cut & tag for CIP at D20 with or without BrdU or IdU.

Therefore, we believe that BrdU/IdU induces DNA damage by incorporating into the genome, and recruits ATM to the site of damage. Subsequently, ATM upregulates the acetylation levels of gene loci through a series of phosphorylation and acetylation cascades, thereby altering chromatin accessibility and activating gene expression.

Combined with the results of RNA-seq, ATAC-seq, motif, γ-H2AX, H3K9ac, and H3K27ac analysis, we found that *Gata4* has a high degree of specificity under the treatment of BrdU/IdU (Figure S2A,2D,2F,3H and 4D). Therefore, GATA4 may be one of the specific downstream factors of BrdU/IdU involved in somatic cell reprogramming. Further Cut & tag analysis showed that the binding sites of GATA4 were significantly increased under the treatment of BrdU/IdU, and the sites enriched with GATA4 were accompanied by higher chromatin accessibility (Figure 4G), including pluripotency gene loci such as *Oct4*, *Sall4* as illustrated in Figure 4H. This indicates that GATA4 can respond to DDR, thereby promoting somatic cell reprogramming.

### BrdU/IdU creates a more open hypomethylated environment for the transformation of cell fate

BrdU and IdU have been demonstrated to play a crucial role in chemical reprogramming, exhibiting two distinct stages of action. Hence we postulate that BrdU not only activates intermediate XEN genes such as *Gata4*, but also potentially induces DNA demethylation, where DNA methylation is a major regulatory factor that limits GATA4 function(*10*). To confirm this, we measured the methylation levels of GATA4 downstream gene *Aqp8* and XEN gene *Pth1r.* We observed a significant decrease in DNA methylation levels with BrdU/IdU treatment (Figure 5A), indicating that BrdU/IdU not only activates gene expression but also induces DNA demethylation, thus creating a conducive environment for pluripotency network activation.

**Fig. 5.**
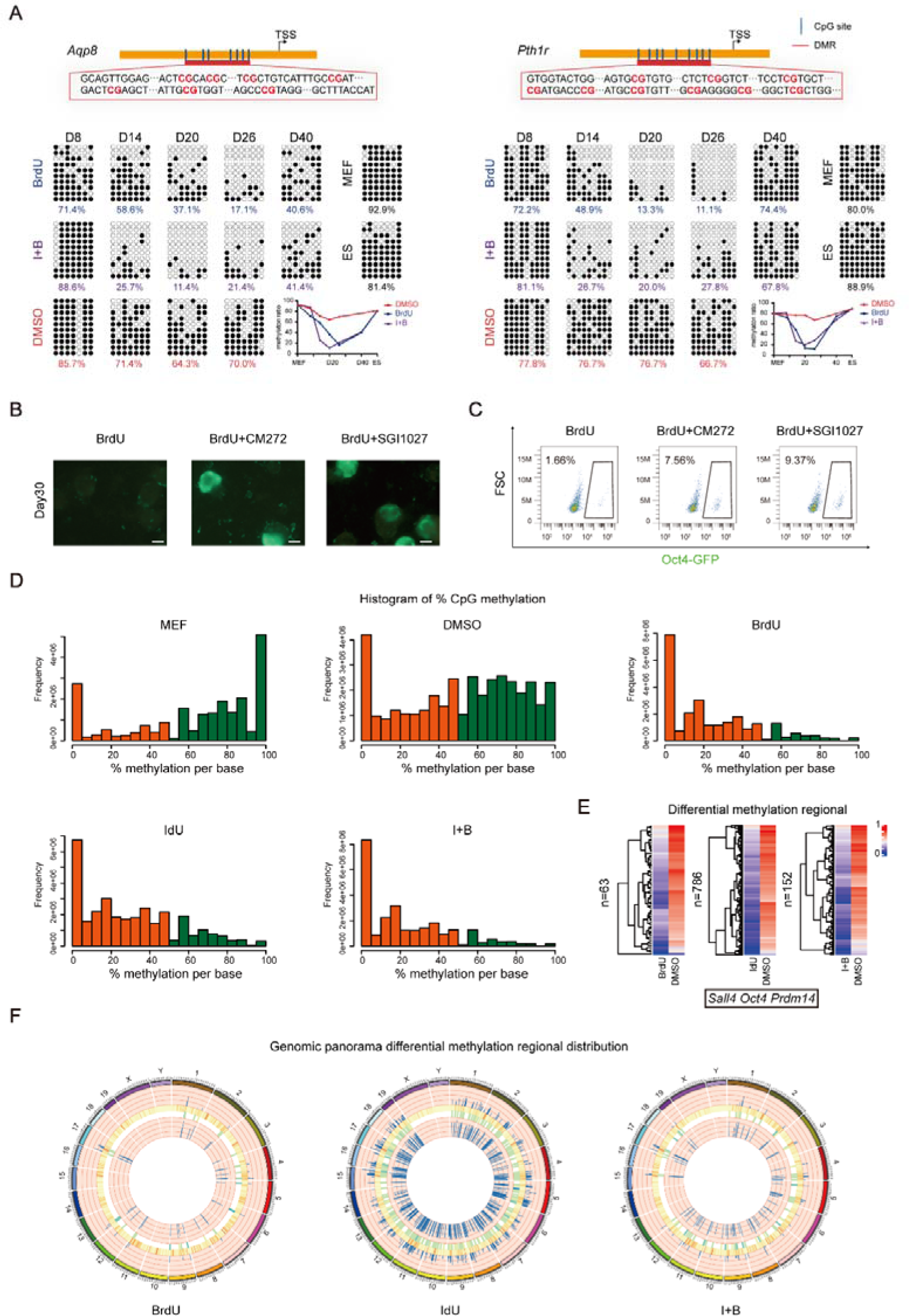
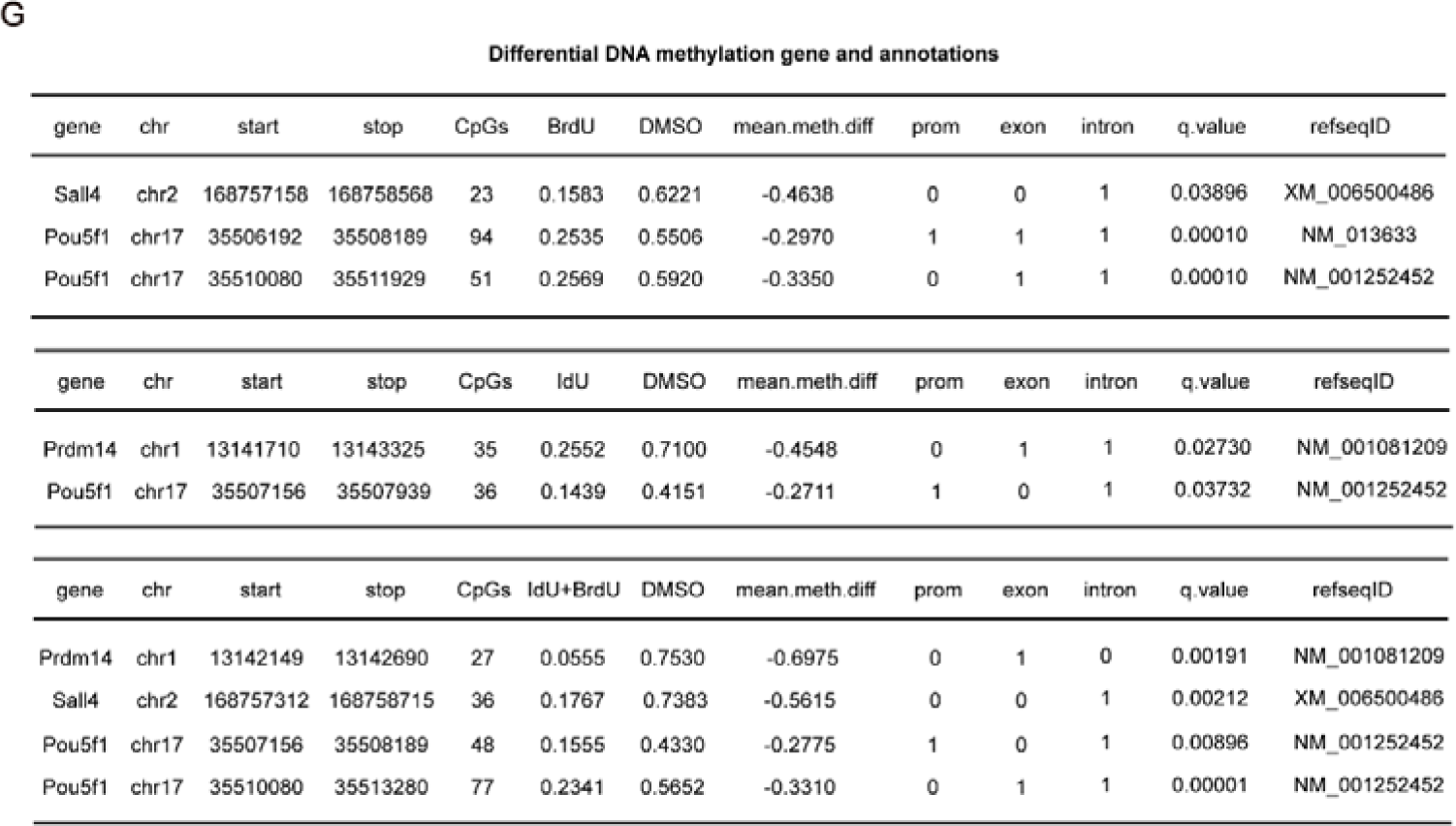
BrdU/IdU creates a more open hypomethylated environment for the transformation of cell fate. (**A**) The methylation patterns of Aqp8 and Pthr1 when treated with or without BrdU or IdU. (**B**)Morphological changes at D30 during induction of CiPSCs treated with DNMT inhibitor CM272 and SGI1027. (**C**) FSC analysis of Oct4-GFP colonies generated under different treatment conditions. (**D**) Histogram of percentage CpG methylation generated under different treatment conditions. (**E**) Methylation differential analysis between treated and untreated cells with IdU/BrdU. (**F**) Genomic panorama differential methylation regional distribution. (**G**) Differential DNA methylation gene and annotations.

We also integrated the aspect of DNA damage repair and hypothesized that BrdU and IdU act through double-stranded break excision and new chain synthesis during the process of DNA damage repair. Since new chain synthesis occurs without DNA methylation marks, it leads to DNA demethylation. To validate this hypothesis, we inhibited DNMT using CM272 and SGI1027 to prevent the re-methylation of new chains. The results indicated that this approach significantly accelerated the process of chemical reprogramming (Figures 5B and C), and this effect was dependent on the presence of BrdU. To summarize, as the degree of BrdU damage increases, the frequency of double-stranded breakage and synthesis also increases, which leads to DNA demethylation, thereby creating a more open hypomethylated environment for the transformation of cell fate.

In the global genome methylation analysis using Gm-seq, we found that the methylation levels of CpG sites after treatment with BrdU, IdU, and I+B were mostly concentrated below 50%, while the distribution was more even in the untreated (DMSO) group, and in MEF it was biased towards above 50% (Figure 5D). The average percentage of CpG sites with a methylated C base was below 30% after treatment with BrdU, IdU, and IB, 50% in the DMSO group, and 60% in MEF (Figure S4A). Furthermore, we used a circos plot to show the distribution of methylation density on chromosomes. The size of each bin was 1,000,000, and the number of methylated Cytosine in each bin with different sequence environments (CpG, CHG, CHH) was counted. The distribution density of CpG sites in different sequence environments throughout the genome showed that after treatment with BrdU/IdU, a significant global hypomethylation state was observed (Figure S4B).

We performed methylation differential analysis between treated and untreated cells with thymidine analogs. Interestingly, we found that the pluripotency related gene *Sall4*, *Oct4*, and *Prdm14* loci exhibited significant hypomethylation after thymidine analog treatment (Figure 5E,G). This further confirms that BrdU/IdU creates a more open hypomethylated environment for the transformation of cell fate. In addition, we found that IdU exhibited a more widespread differential methylation distribution, followed by I+B, and BrdU exhibited the least, which may be related to its atomic size (Figure 5F).

### BrdU/IdU can participate in cell fate regulation with negligible mutations

As the substitute of thymidine, the safety of BrdU has always been a concern. Therefore, we performed mutation analysis on the samples treated with or without thymidine analogs and MEFs in our system, including SNP (Single Nucleotide Polymorphism), InDel (insertion-deletion), CNV (Copy Number Variant), and SV (Structural variation). We found that the mutation level of BrdU/IdU-treated samples was almost the same as that of MEFs (Figure 6A-E), while no treatment with thymidine analogs led to more mutations. In addition, we found that IdU had some insertion mutations. Overall, this suggests that the rational use of BrdU/IdU can participate in cell fate regulation with negligible mutations.

**Fig. 6.**
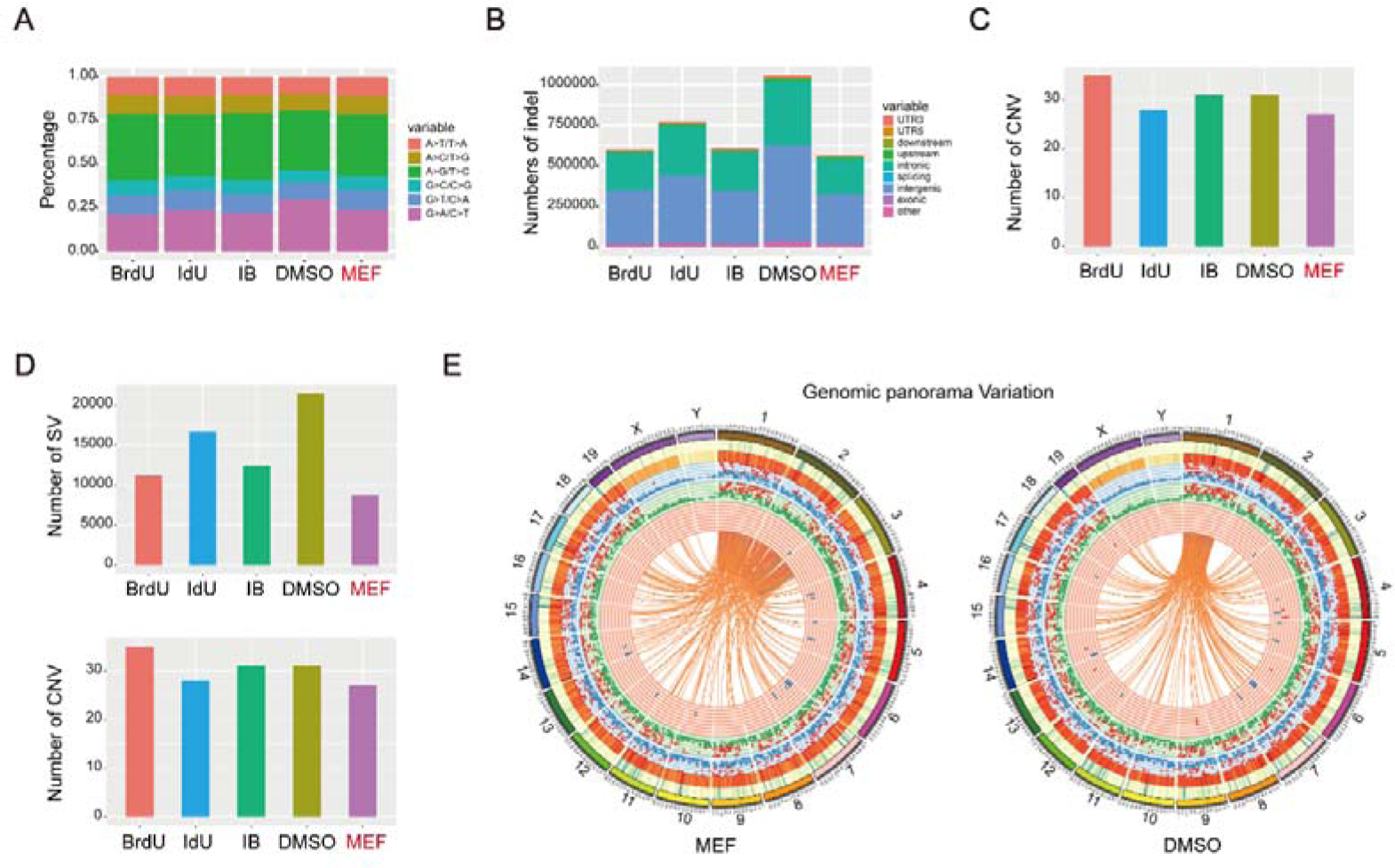
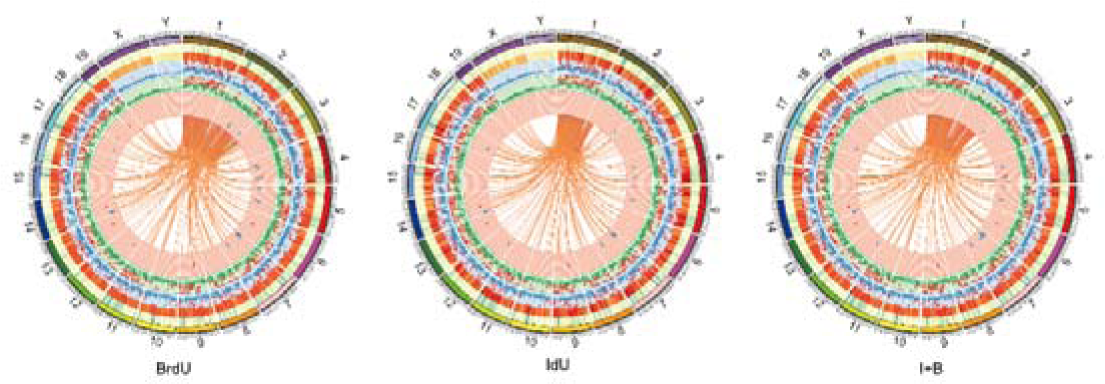
BrdU/IdU can participate in cell fate regulation with negligible mutations. (**A**) Mutation analysis-SNP (Single Nucleotide Polymorphism) on the samples treated with or without BrdU/IdU and MEFs. (**B**) InDel (insertion-deletion) analysis. (**C**) CNV (Copy Number Variant) analysis. (**D**) SV (Structural variation) analysis. (**E**) Genomic panorama variation analysis.

### BrdU and IdU have functions in various reprogramming systems

To investigate whether the effects of BrdU and IdU are applicable in other reprogramming systems, we found that BrdU significantly improves reprogramming efficiency in the OKS system (Figure S5A-D, Figure 7AB). Additionally, we found that BrdU can substitute for OCT4 and play a critical role in the KS system (Figure S5E,F,G, Figure 7A,B). Incorporation of BrdU not only improves reprogramming efficiency, but also accelerates reprogramming speed (Figure 7C). Interestingly, by comparing the effects of BrdU on OKS, KS, and CIP (Figure 7A,B), we found that their dependence on BrdU treatment time gradually increases. Specially, in OKS system, GFP^+^ colonies reach their peak when BrdU is treated for 2 days. In KS system, GFP^+^ colonies reach their peak when BrdU is treated for 6 days. Furthermore, in CIP system, 22 days of BrdU treatment is required to reach maximum number of GFP^+^ colonies, as illustrated in Figure 7B. This suggests that different degrees of DNA damage may be required to overcome various barriers to reprogramming.

**Fig. 7.**
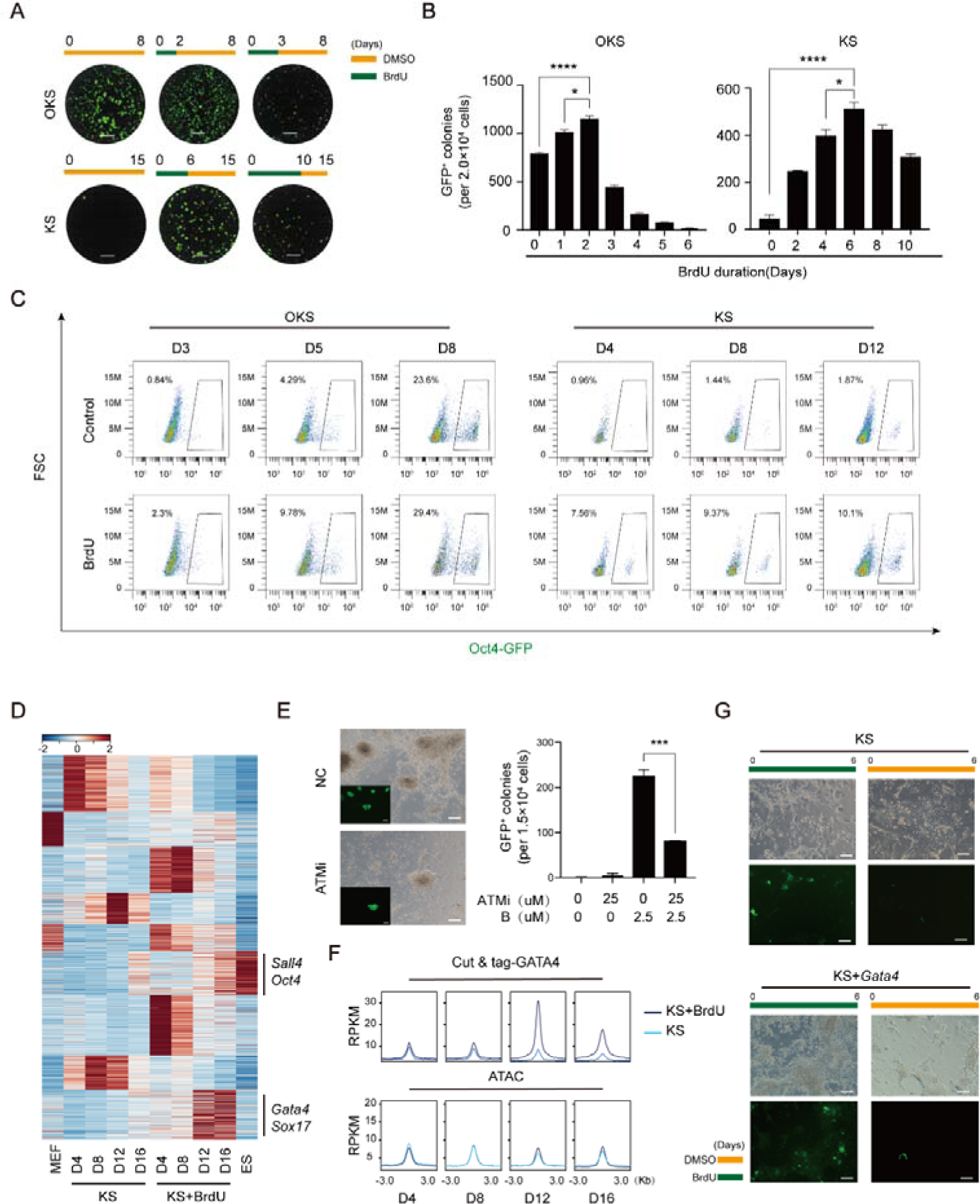
BrdU and IdU have functions in various reprogramming systems. (**A**) Images of GFP+ colonies taken by fluorescence microscope in situ. Scale bar 5mm. (**B**) Number of Oct4-GFP colonies generated under indicated conditions. (**C**) FSC analysis of Oct4-GFP colonies generated under different treatment conditions. (**D**) Heatmap of RNA-seq for KS at D4,8,12 and 16 with or without BrdU or IdU. (**E**) Morphological changes during induction of KS treated with ATM inhibitor and corresponding statistical analysis. Scale bar 50µm. (**F**) GATA4 Cut & tag analysis and ATAC-seq analysis for CIP at D4,8,12 and 16 under different treatment conditions. (**G**) Morphological changes during induction of KS when overexpressing Gata4 in the system. Scale bar 50µm

We discovered that the addition of BrdU in the KS reprogramming system can specifically activate *Gata4*, *Sox17*, and certain developmental stem cell pluripotency regulating genes, as shown in Figure 7D, Additionally, we observed that the most significant difference occurred at D12, which was precisely the time point when genes such as *Gata4* were highly expressed. We found that inhibiting the ATM signaling pathway with an ATM inhibitor also prevented BrdU from functioning in KS system (Figure 7E). By using GATA4 Cut & tag, we discovered that GATA4-enriched peaks were accompanied by ATAC opening (Figure 7F). By overexpressing *Gata4* in the KS system under the presence of BrdU, the reprogramming process can be accelerated and the cell state can be improved (Figure 7G). Remarkably, our results bear a resemblance to the phenomenon observed in chemical reprogramming.

## Discussion

In summary, we found that the use of the thymidine analogs BrdU/IdU causes epigenetic reshaping. This potentiates cells to enter a plastic state and accelerates the fate transition of cells by activating DNA damage repair and causing significant H3K27ac, H3K9ac and DNA demethylation. Additionally, the study discovered that the XEN state can be specifically activated after BrdU/IdU treatment. This suggest that XEN state may be a selective response behavior of cells to DNA damage repair, providing a shortcut for cells to overcome reprogramming barriers, and that different degree of DNA damage may be required to overcome various barriers to reprogramming. Furthermore, rational doesage of BrdU/IdU can participate in cell fate transition with negligible mutantations.

In our study, we observed that *Apex1*, a gene related to BER pathway, was significantly upregulated in the early stages of reprogramming after treatment with BrdU/IdU, but gradually decreased with longer exposure. The HRR related genes *Brca1* and *Brca2* were upregulated as *Apex1* expression decreased (Figure S3C). This suggests a shift from BER to HRR for DNA damage repair, with a more intense HRR pathway causing an overall epigenetic reshaping of cells and enabling them to overcome greater barriers. However, it is worth to investigated the roles of other DNA repair pathways, such as MMR, NHRJ and NER, in the process of cellular fate transition. Additionally, not all of thymidine analogs are equally effective in somatic reprogramming, and their specific mechanisms of action remain to be fully explored. Interestingly, although BrdU and IdU are structurally very similar, with only minor differences in their chemical structure, the difference in atomic size may lead to a more pronounced impact of IdU on the entire genome. This difference in impact may also explain why other thymidine analogs are not effective.

It is worth to consider that the XEN-like intermediate state is not unique to chemical reprogramming, as cells may require multiple intermediate states to reach their final fate during the process of fate transition. Reprogramming barriers may limit the process of fate transition, and thus cells may choose different intermediate states as shortcuts. *Gata4* and *Sox17*, among others, are representative of XEN states that have been selected as transitional states that facilitate cell fate transition because they promote both dedifferentiation and redifferentiation of cells, and play essential roles in embryonic development and cell regeneration. Moreover, DNA damage repair may also drive cells into a plastic state. When DNA damage occurs, cells need to repair the damage to maintain genome integrity. DNA repair processes may lead to reprogramming events, resulting in cells entering a plastic state and providing an alternative pathway for cell fate transition.

## Acknowledgements

This work was supported by grants from The National Key Research and Development Program of China (2018YFE0204800,XX); National Natural Science Foundation of China (32022019, 31970681, 32100594, 32200695); Grant No:QYZDJ-SSW-SMC009; Science and Technology Planning Project of Guangdong Province No. 2017B030314056, 2019A1515011024; Key Research & Development Program of Guangzhou Regenerative Medicine and Health Guangdong Laboratory(2018GZR110104003). Science and Technology Projects in Guangzhou, China. Grant No. (202201010602, 202201010510); China Postdoctoral Science Foundation (2022000312); Science and Technology Planning Project of Guangdong Province. Grant No. (2020B1212060052, 2022B1212010010).

## Author contributions

J.L. and C.L. initiated the project and designed the experiments. C.L. performed the chemical reprogramming experiments. C.L conducted and performed ATAC-seq, RNA-seq, Cut & tag library construction with X.X. S.C and Chen Li preformed bioinformatics analysis. X.X.performed OKS and KS reprogramming experiments with C.L. X.X performed bisulfite sequencing PCR. C.L performed GM-seq with Gene plus company. J.L. D.P. and H.W supervised the whole study. H.W supervised the whole study and wrote the manuscript. J.L conceived the whole study and approved the final version.

## Competing interests

The authors declare no competing financial interests.

## Materials and methods

### Mice

Beijing Vital River Laboratory supplied the 129Sv/Jae and ICR mice, while The Jackson Laboratory provided the Oct4-GFP(OG2) transgenic allele-carrying mice (CBA/CaJ X C57BL/6J). All animal experiments were carried out in compliance with the Animal Protection Guidelines of Guangzhou Institutes of Biomedicine and Health, located in Guangzhou, China and affiliated with the Chinese Academy of Sciences.

### Cell culture

#### Mouse embryonic fibroblasts (MEFs)

To isolate Mouse Embryonic Fibroblasts (MEFs), E13.5 embryos were dissected and all internal organs, head, limbs, and tails were removed and discarded. The remaining tissue was sliced into small pieces and dissociated with a digestive solution (consisting of 0.25% trypsin and 0.05% trypsin in a 1:1 ratio from GIBCO) for 15 minutes at 37C to obtain a single-cell suspension. The cells from each embryo were then plated onto a 6-cm culture dish coated with 0.1% gelatin and cultured in DMEM (Hyclone) supplemented with 10% FBS (GIBCO), 1% GlutaMAX (GIBCO), and 1% NEAA (GIBCO), referred to as fibroblast medium.

mESCs

The feerder-free and serum-free ESC medium used for maintaining mESCs and CiPSCs consisted of DMEM supplemented with N2 (100X), B27 (50X), LIF, 1% GlutMax, 1% NEAA, 0.1 mM beta-mercaptoethanol, and 3 µM CHIR99021 and 1 µM PD0325901. Additionally, the cell lines were tested negative for mycoplasma using Lonza’s Kit (LT07-318).

Culture medium preparation Chemical reprogramming stage 1:

iCD1 medium (Chen et al., 2011a) comprises Vitamin C (50 mg/ml), bFGF (10 ng/ml), and CHIR99021 (3 mM), along with several small molecules, including Brdu (10 µM), RepSox (5 mM), FSK (10 mM), VPA (0.1 mM), AM580 (0.05 mM), EPZ5676 (5 mM), DZNeP (0.05 mM), SGC0946 (5 mM), BMP4 (10 ng/ml), and Capmatinib (5µM).

Chemical reprogramming stage 2:

DMEM (high glucose, Hyclone), 1% N2 (GIBCO), 2% B27 (GIBCO), 1% GlutMax (GIBCO), 1% NEAA (GIBCO), 1% sodium pyruvate (GIBCO) 3 mM CHIR99021, 1 mM PD0325901 and 1000 U LIF.

KS reprogramming stage 1:

iCD1 medium, Repsox(5 μM), FSK(10 μM), SGC0946(1 μM), GSK-LSD1(1 μM) and BrdU(2.5 μM).

KS reprogramming stage 2:

iCD1 medium, Repsox(5 μM), FSK(10 μM), SGC0946(1 μM), GSK-LSD1(1 μM).

KS reprogramming stage 3:

DMEM (high glucose, Hyclone), 1% N2 (GIBCO), 2% B27 (GIBCO), 1% GlutMax (GIBCO), 1% NEAA (GIBCO), 1% sodium pyruvate (GIBCO) 3 mM CHIR99021, 1 mM PD0325901 and 1000 U LIF.

### Chemical induction of iPSCs from mouse fibroblasts

The 12-well plates were pre-coated with 0.1% gelatin and seeded with MEFs at a density of 20,000 cells per well in 10% fibroblast medium. The following day, Stage 1 chemical reprogramming medium was added and refreshed daily. After day 22, the medium was changed to Stage 2 chemical reprogramming medium. On day 40, the *Oct4*-GFP colonies were either counted or detected via FACS.

*Klf4* and *Sox2* mediated MEF reprogramming

Plasmids carrying murine Klf4, Sox2 cDNA were purchased from Addgene, and the fragments were then ligated to the PMX vector to obtain a recombinant plasmid. PlatE cells were transfected with individual plasmids with the Ca3(PO4)2-method. After 48 hours of transfection, the virus supernatant was collected and used to infect cells. Approximately 15,000 cells were seeded per well in 24-well plates containing DMEM medium supplemented with 10% FBS, and were cultured for 24-12 hours prior to MEF infection. The Klf4 and Sox2 virus solution was filtered using a 0.45 µm filter, and then 8 µg/ml of polybrene was added for cell infection. The cells were subjected to a second infection after 24 hours. After 48 hours of MEF cell infection, the cells were transferred to induction medium, with the day of medium change being designated as day 0. KS reprogramming stage 1 medium was used for the first 6 days, stage 2 medium was used from day 6 to day 10, and stage 3 medium was used from day 10 onwards. The medium was changed every 24 hours.

### Immunofluorescence staining

The coverslip-grown cells were washed twice with PBS and fixed using 4% paraformaldehyde (PFA) at room temperature for 30 minutes. Next, the samples were treated with 1N HCl on ice for 10 minutes, followed by 2N HCl at room temperature for 10 minutes and then 2N HCl at 37°C for 20 minutes. After washing thrice with PBS for 5 minutes each, the cells were permeabilized with 0.1% Triton X-100 for 30 minutes. Subsequently, the cells were blocked with 3% BSA for an hour and washed thrice with PBS for 5 minutes each while shaking. The cells were then incubated overnight at 4°C with primary antibodies, diluted in 3% BSA (1:250) in PBS. After four washes with PBS for 5 minutes each while shaking, the cells were incubated for an hour at room temperature with secondary antibodies, diluted in 3% BSA (1:200) in PBS. Later, the cells were incubated in DAPI for 1 minute, washed twice with PBS, and finally, the coverslips were mounted on the slides for observation under a confocal microscope (Andor Dragonfly 200). The following antibodies were used: anti-BrdU (Sigma), anti-γ-H2AX (Abcam), secondary antibodies Alexa Fluor 568 goat anti-mouse IgG (Invitrogen) and Alexa Fluor 488 goat anti-mouse IgG (Invitrogen).

### Comet assay

During the chemical reprogramming process, the control and experimental group cells at D14, D16, D24, and D30 were digested with 0.25% trypsin into single cells, and resuspended in PBS at a concentration of 10,000 cells/ml. The Oxiselect Comet Assay Kit (Cell-Biolabs) was used as a reference for the experiment. The cell suspension was mixed with agarose, and then incubated in the dark at 4℃ for 15 minutes. The mixture was placed on a slide and immersed in lysis buffer, then incubated in the dark at 4℃ for 60 minutes. The slide was transferred to an alkaline solution, and incubated in the dark at 4℃ for 30 minutes. The slide was then placed into an electrophoresis chamber, and alkaline electrophoresis buffer was added under the conditions of 300 mA and 15 minutes. After electrophoresis, the slide was transferred to water and soaked for 2 minutes. Then it was transferred to 70% ethanol and soaked for 5 minutes. The slide was placed in a 37℃ oven overnight. Vista Green DNA dye was added for 10 minutes, and the cells were observed and recorded using an inverted fluorescence microscope (ZEISS). The software cometscore was used to analyze the tail moment of the cells.

### FACS analysis

The cells were rinsed with PBS and then digested with 0.25% trypsin at 37°C for 5 minutes. The digestion was stopped by adding 10% FBS DMEM medium. After filtering with a sieve and centrifuging at 250 g for 5 minutes, the supernatant was discarded and the cells were resuspended in PBS at a concentration of 100,000 to 1,000,000 cells/ml. The percentage of positive cells was detected using a BD Accuri™ C6 Plus flow cytometer. The flow cytometry data was analyzed using FlowJo7.6.1.

### Bisulfite genomic sequencing

Extract DNA template. After centrifuging the cells, add 600 μl of Nuclei Lysis Solution (Promega) and mix by pipetting. Add 200 μl of Protein Precipitation Solution (Promega) and mix by pipetting. Let it sit on ice for 10 minutes. Centrifuge at 12,000 rpm for 10 minutes. Take the supernatant, add 600 μl of isopropanol, and let it sit on ice for 10 minutes. Centrifuge at 12,000 rpm for 5 minutes. Discard the supernatant, add 600 μl of 70% ethanol. Heat in a metal bath at 60°C for 5 minutes. Add RNAase-free water and heat in a metal bath at 60°C for 30 minutes. Take 2 μg of template DNA and bisulfite treat according to the EpiTect Bisulfite Kit (QIAGEN) manual. Amplify the Aqp8 and Pth1r promoter regions by PCR. Clone and sequence using the pMD18-T vector (TaKara).

### RNA-seq and data analysis

Cells were treated with TRIzol to extract total RNA, which was then converted into cDNA using ReverTra Ace (Toyobo) and oligo-dT (Takara). The resulting cDNAs were analyzed using Premix Ex Taq (Takara) in qPCR experiments. For library construction, the TruSeq RNA Sample Prep Kit (RS-122-2001, Illumina) was employed, and RNA-seq was performed using the Miseq Reagent Kit V2 (MS-102-2001, Illumina).

The original sequencing data was quality-controlled using FASTQC, and low-quality bases and sequencing adapters were removed using trim_galore. HISAT2(*11*) was used to align the filtered clean reads with the mouse mm10 reference genome. Samtools(*12*) was used to filter out unaligned or unpaired sequencing fragments and obtain the bam file. featureCounts(*13*) software was used to quantify gene expression levels. EdgeR was used to identify differentially expressed genes. ClusterProfiler(*14*) was used to perform GO or KEGG functional enrichment analysis to determine the molecular functions of the differentially expressed genes.

### ATAC-seq

The ATAC-seq procedure was carried out following the previously published protocols(*15,16*) and TruePrep DNA Library Prep Kit V2 for Illumina. Briefly, 50,000 cells were washed with 50 mL of cold PBS and suspended in 50 mL of lysis buffer containing 10 mM Tris-HCl pH 7.4, 10 mM NaCl, 3 mM MgCl2, and 0.2% (v/v) IGEPAL CA-630. The cell suspension was centrifuged at 500 g for 10 minutes at 4°C, followed by the addition of 50 mL of transposition reaction mix from the TruePrep DNA Library Prep Kit V2 for Illumina. The resulting samples were subjected to PCR amplification and incubated at 37°C for 30 minutes. VAHTS DNA Clean Beads (Vazyme #N411) were used for purification and recovery of the DNA fragments.The purified product was amplified. VAHTS DNA Clean Beads were used for further purify and separate 200-700bp fragments. The ATAC library was finally sequenced on a NextSeq 500, using a NextSeq 500 High Output Kit v2 (150 cycles) (FC-404-2002, Illumina), following the manufacturer’s instructions.

### Cut&tag

According to the protocol of the Hyperactive universal Cut&Tag assay Kit for Illumina, library construction was performed. 100,000 cells were collected and incubated with processed ConA beads. The sample was incubated with primary antibody overnight at 4℃, followed by incubation with secondary antibody at room temperature for 1 hour. The sample was then incubated with transposase pA/G-Tnp at room temperature for 1 hour, and the DNA was fragmented for 1 hour before extraction. After amplifying DNA fragments according to concentration, VAHTS DNA Clean Beads (Vazyme #N411) were used for purification and recovery of the DNA fragments.

### ATAC-seq and Cut&tag bioinformatics analysis

The original sequencing data was quality-controlled using FASTQC, and low-quality bases and sequencing adapter sequences were removed using trim_galore. Bowtie2(*17*) was used to align high-quality sequencing fragments with the mouse mm10 reference genome. Samtools was used to retain only paired and uniquely aligned sequences, exclude mitochondrial sequence fragments, and obtain the bam file. Picard(http://broadinstitute.github.io/picard/) was used to remove duplicate fragments caused by PCR amplification during library preparation. Deeptools(*18*) was used to convert the bam file to a bw file, and the results were visualized using IGV(https://igv.org/). The peak calling software MACS2(*19*) was used to detect enrichment regions of open chromatin or DNA fragments. HOMER was used to identify transcription factor binding motifs enriched in chromatin regions. CHIPseeker(*20*) was used to annotate genomic features (such as genes, promoters, enhancers, and transcription factor binding sites) of peaks intervals. Bedtools(*21*) was used to analyze differential binding sites, and deeptools was used to generate signal matrix files and visualize the results. Other analyses were performed using glbase(*22*).

### GM-seq

The Gm-seq procedure was carried out following the previously published protocols in cooperation with Gene plus company(*23*). We utilized Hieff NGS® Ultima Pro DNA Library Prep Kitfor Illumina (Yeason, cat. on. 12201ES96) for end repair, A-tailing, and adaptor ligation. After magnetic beads were purified, 5-methylcytosine (5mC) and 5-hydroxymethylcytosine (5hmC) were oxidized to 5-acylcytosine (5fC) or 5-carboxyl cytosine (5caC) by TET enzyme, then 5fC or 5caC was treated with pyridine borane and reduced to dihydrouracil (DHU). Finally, PCR amplification is performed and barcode is introduced to obtain the sequencing library. After going through this series of processes the methylated cytosine would be identified as thymine (T) by MGI T7 sequencing platform. The samples for whole genome sequencing directly were also going through the same procedures that were used for GM-seq but without oxidation and pyridine borane reduction preparation. Whole genome analyses were performed by the sequencing library we constructed above.

### GM-seq bioinformatics analysis

The raw fastq and bam files were evaluated using BWA and SAMtools software. Alignment and analysis of DNA sequencing data were performed utilizing asTair tools in automated mode from raw fastq files to finalized genotyping data. This pipeline utilized GATK tools and ExomeDepth tools to perform base quality score, genotype calling and variant annotation, including SNP and Indel. And ichorCNA were employed to analyze genome-wide CNV and calculate insert size. Sequencing data were analyzed by three computational pipelines in order to understand the reliability of the detection of fungi, bacteria, and viruses: Kraken, Kaiju and SNAP. BLAST was used to compare the sequencing results with the high-risk microorganisms. For methylation analyses, diagrams for CpG coverage were generated using methylKit 1.4.0. The GC bias plot was generated using Picard’s CollectGCBiasMetrics. Correlation analysis was performed using methylKit version 1.4.0 with default settings except for a minimum coverage threshold of one read.

### Quantification And Statistical Analysis

Sample size was not predetermined using statistical methods, and the experiments were not randomized. The investigators were not blinded during the experiment or outcome assessment. The data was presented as mean±s.d. Statistical analysis was conducted using either two-tailed unpaired student’s t-test or one-way ANOVA with GraphPad Prism 6, and a P-Value of <0.05 was considered statistically significant. The level of significance was indicated as ∗P < 0.05, ∗∗P < 0.01, ∗∗∗P < 0.001 and ∗∗∗∗P < 0.001. The relevant figure legends provided information on the statistical test, precise P-Value, exact sample sizes, and independent experiments.

## Supplementary Figure

**Figure S1:**
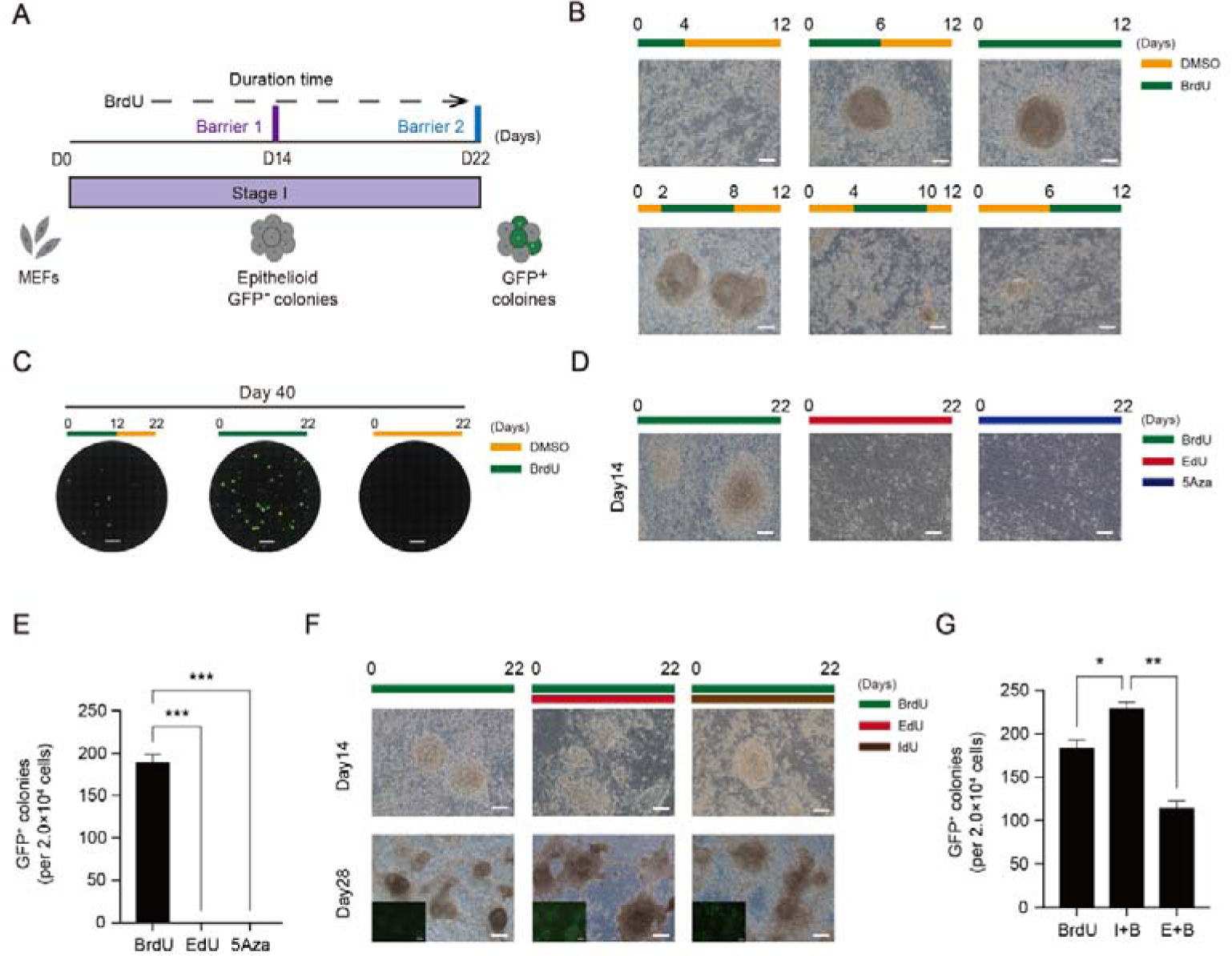
A. Schematic diagram of the induction of stage1 B. Morphological changes at barriers1 under indicated conditions. Scale bar 100µm C. Images of GFP+ colonies taken by fluorescence microscope in situ. Scale bar,5mm. D. Morphological changes at barriers1 treated with BrdU, EdU or 5Aza. E. Number of *Oct4*-GFP+CiPSC colonies generated under BrdU,EdU or 5Aza. F. Morphological changes at distinct time points during induction of CiPSCs treated with BrdU, I+B(IdU+BrdU) or B+E(BrdU+EdU). Scale bar 100µm G. Number of *Oct4*-GFP+CiPSC colonies generated under BrdU,I+B or B+E.

**Figure S2:**
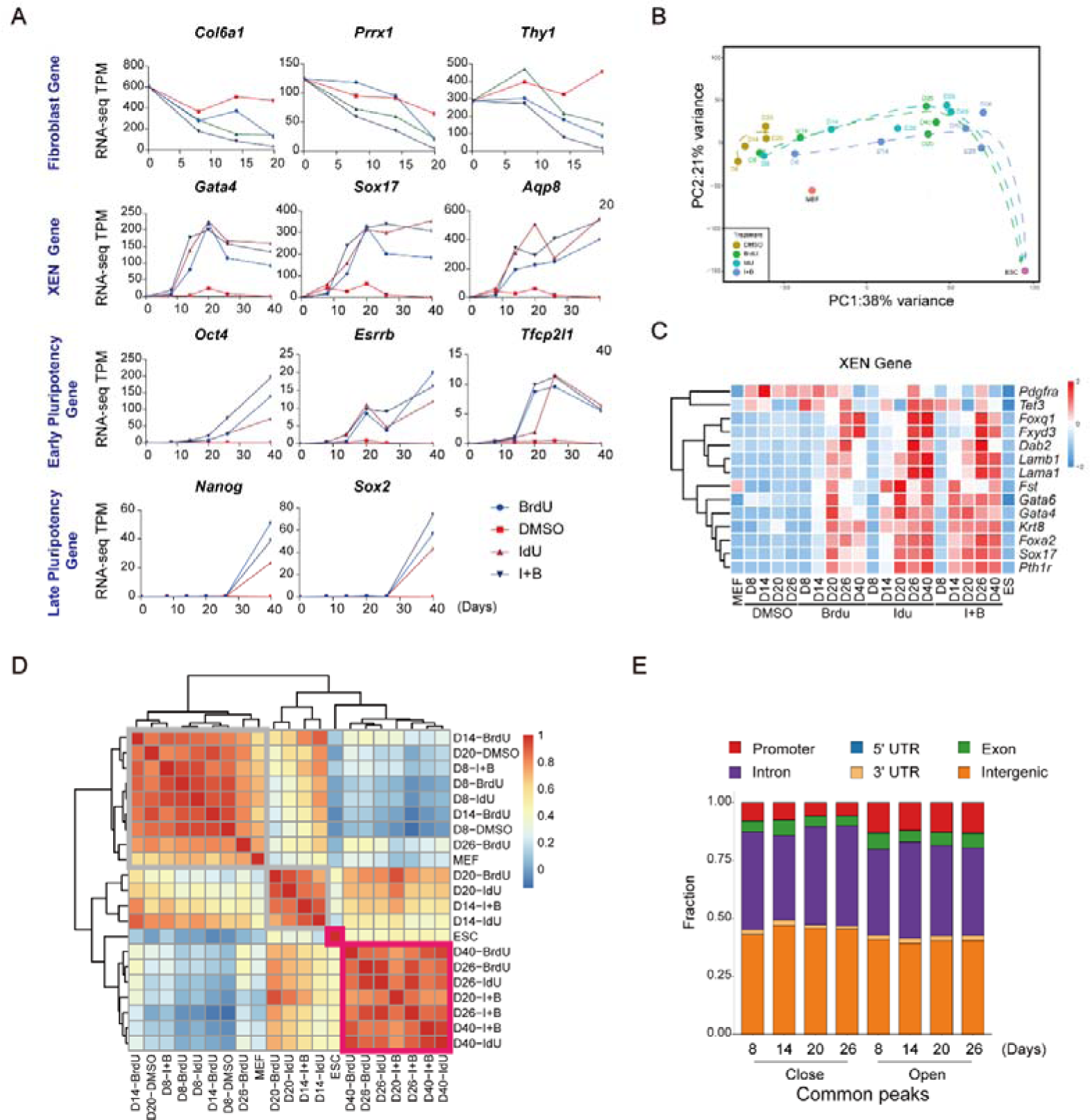
A. Representative fibroblast genes, XEN genes and pluripotency genes peaks from RNA-seq for CIP at D8,14,20 and 26 with or without BrdU or IdU. B. PCA analysis for RNA-seq data from Cip under indicated conditions C. Heatmap of XEN genes related RNA-seq for CIP at D8,14,20 and 26 with or without BrdU or IdU. D. The R2 correlation coefficient matrix of all versus all RNA-seq datasets as indicated

**Figure S3:**
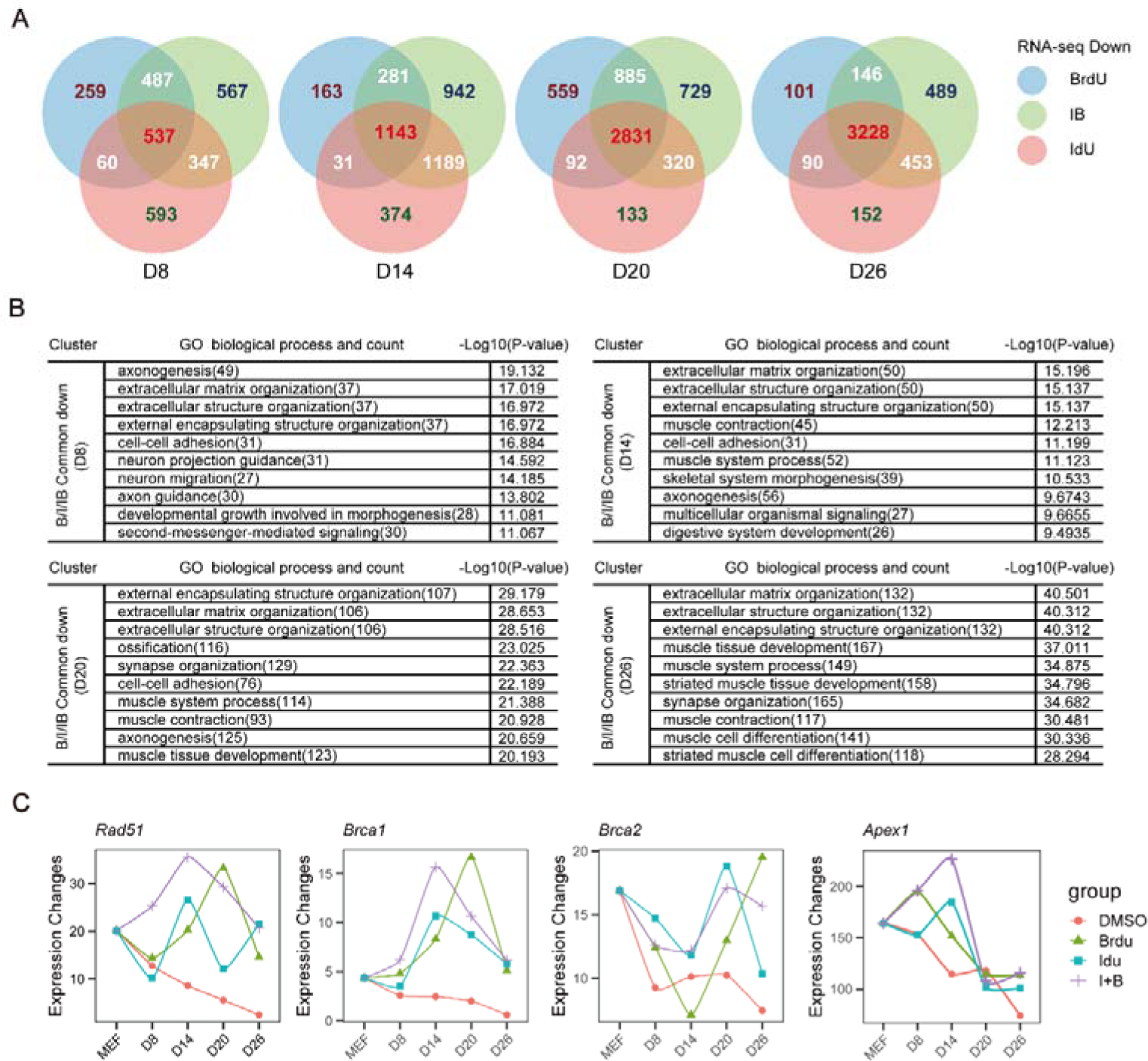
A. Venn plots for overlapping genes in the downregulated groups between BrdU, IdU, and BrdU+IdU versus the control. B. Enriched GO functions in upregulated groups. C. Representative DNA repair genes from RNA-seq for CIP.

**Figure S4:**
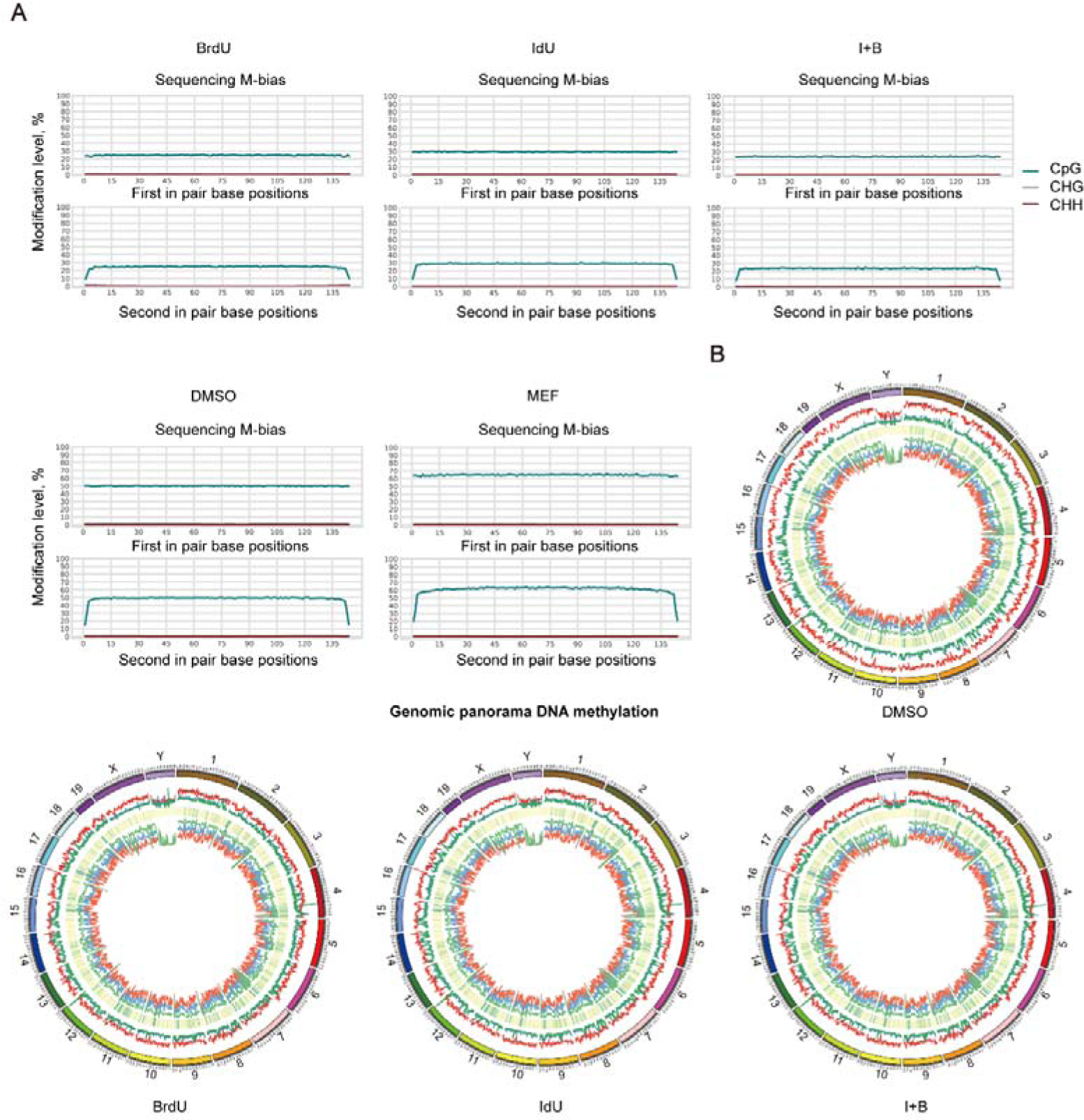
A. M-bias plot analysis B. Genomic panorama DNA methylation.

**Figure S5:**
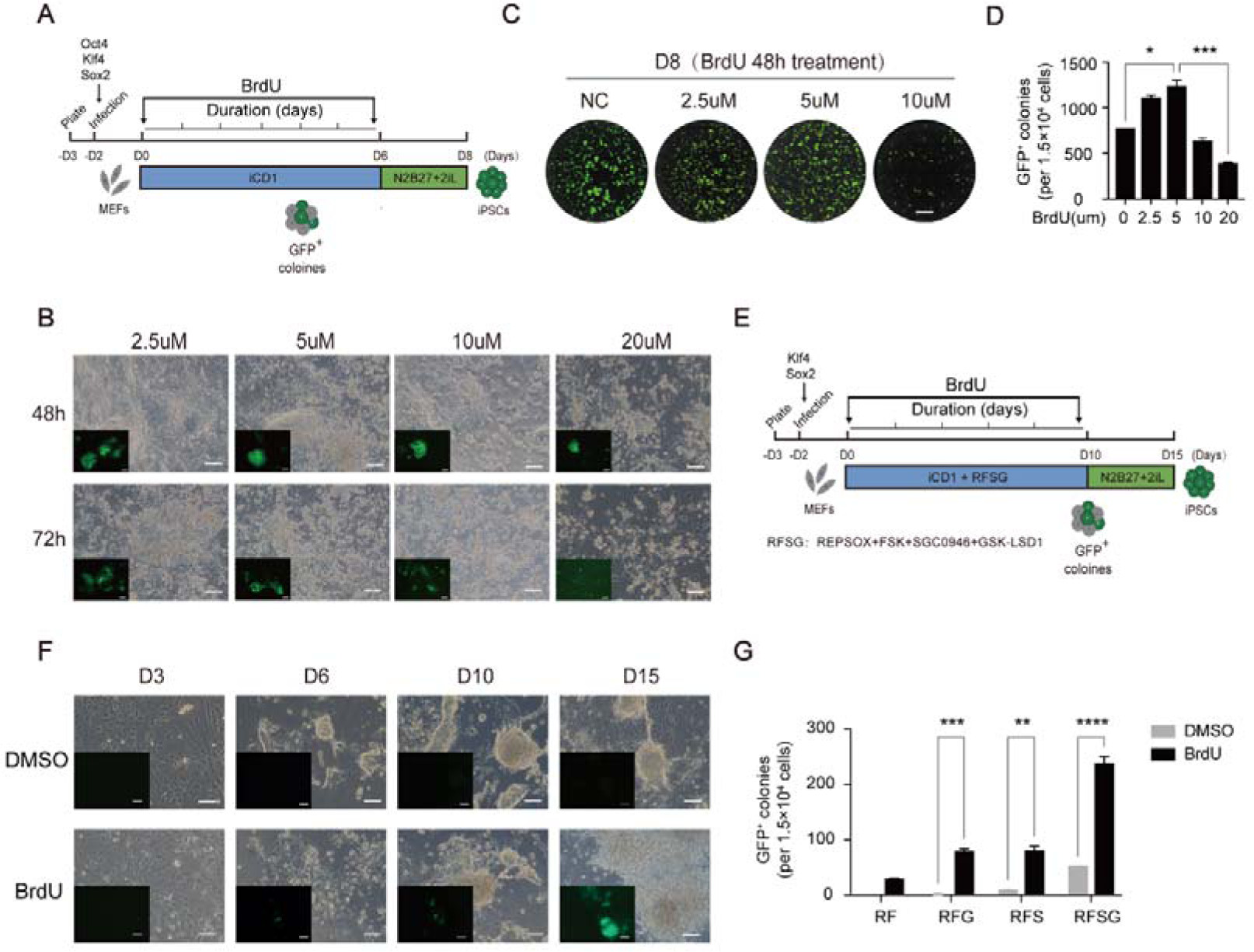
A. Schematic diagram of the induction of iPS from MEFs with OKS B. Images of GFP+ colonies taken by fluorescence microscope in situ. Scale bar,5mm. C. Number of *Oct4*-GFP+CiPSC colonies generated under indicated conditions. D. Morphological changes at OKS treated with different concentrations BrdU E. Schematic diagram of the induction of iPS from MEFs with KS F. Morphological changes at KS treated with or without BrdU G. Number of *Oct4*-GFP+CiPSC colonies generated under indicated conditions. R,Repsox,F,FSK,S,SGC0946,G,GSK-LSD1

